# Adaptation of the autosomal part of the genome on the presence of dioecy

**DOI:** 10.1101/2021.08.05.455242

**Authors:** Jitka Žlůvová, Roman Hobza, Bohuslav Janoušek

## Abstract

We have attempted to answer the question of whether the presence of sex chromosomes in the genome can affect the evolution of the autosomal part of the genome. As a model, we used dioecious plants from the section *Otites* of the genus *Silene*. We have observed a rise in adaptive evolution in the autosomal and pseudoautosomal parts of the genome, which are associated with the evolution of dioecy. This rise is caused neither by the accumulation of sexually antagonistic genes in the pseudoautosomal region nor by the co-evolution of genes acting in mitochondria (in spite of the fact that the dioecy evolved in this case most likely from cytoplasmic male sterility). Thus, this rise in the amount of positively selected codons is most likely caused by the adaptive evolution of genes involved in the specialization of the autosomal part of the genome on the dioecy as described in sex-allocation theory.

## Introduction

Separate sexes are very common in animals, but they also appear in plants (Renner & Müller, 2021). After sex determining gene(s) appear in the genome, many evolutionary processes are started on the sex chromosomes (Charlesworth, 2019). Moreover, many autosomal genes express in a sex-specific manner, and their expression is regulated by the presence of sex-determining gene(s). These sex-specific differences in gene expression lead to the formation of male and female individuals. But can the presence of separate sexes lead to evolutionary changes in the sequences of autosomal genes? Because males invest more in the quantity of their progeny, whereas the goal of the females is to invest in the quality of the progeny, the presence of separate sexes could lead to the adaptation of the autosomal genes to males’ and females’ different breeding strategies.

In animals, separate sexes and sex chromosomes are of very old origin, and thus it could be very difficult to search for signs of the adaptation of the autosomal part of the genome to the presence of separate sexes. On the other hand, many plant genera contain recently evolved dioecious clades. One of the plant genera with young sex chromosomes is the genus *Silene*. The dioecious *Silene* species are especially well-suited to this study because closely related non-dioecious species are known, transcriptomic data for both the dioecious species and their non-dioecious relatives are known, and genetic maps are available in several species (reviewed in (Balounova et al., 2019)). In the genus *Silene*, apart from the dosage compensation studies (Martin et al., 2019; Muyle et al., 2018), adaptation of the transcriptome to the dioecy was also studied from a quantitative point of view (Zemp, Widmer, & Charlesworth, 2018). However, it is not known whether the presence of sex chromosomes in the genome can affect the evolution of the coding regions in the autosomal part of the genome.

## Results and discussion

In this study, we use the currently available RNAseq data obtained in the dioecious section *Otites* to study the influence of the rise of dioecy on the evolution of the sequences that are not sex-linked. Results of the phylogenetic analysis shown in Fig. 1 are mostly in accordance with the results from the previous study that did not include *S. nocturna* and *S. paradoxa*. A minor difference in topology is that the current results support the clade joining two members of the group Cyri (defined previously (Balounova et al., 2019) according to phenotypic markers), while in our previous study, the group Cyri appeared as completely polyphyletic (Balounova et al., 2019). Current dating, based on the estimation of the synonymous divergence of the outgroup (S. nocturna) from the other species, suggests the age of dioecious section *Otites* cca 2.3 million years which is in the range of the previous estimate based on calibration by fossils (1.17–2.60 million years) (Slancarova et al., 2013).

**Figure 1.**
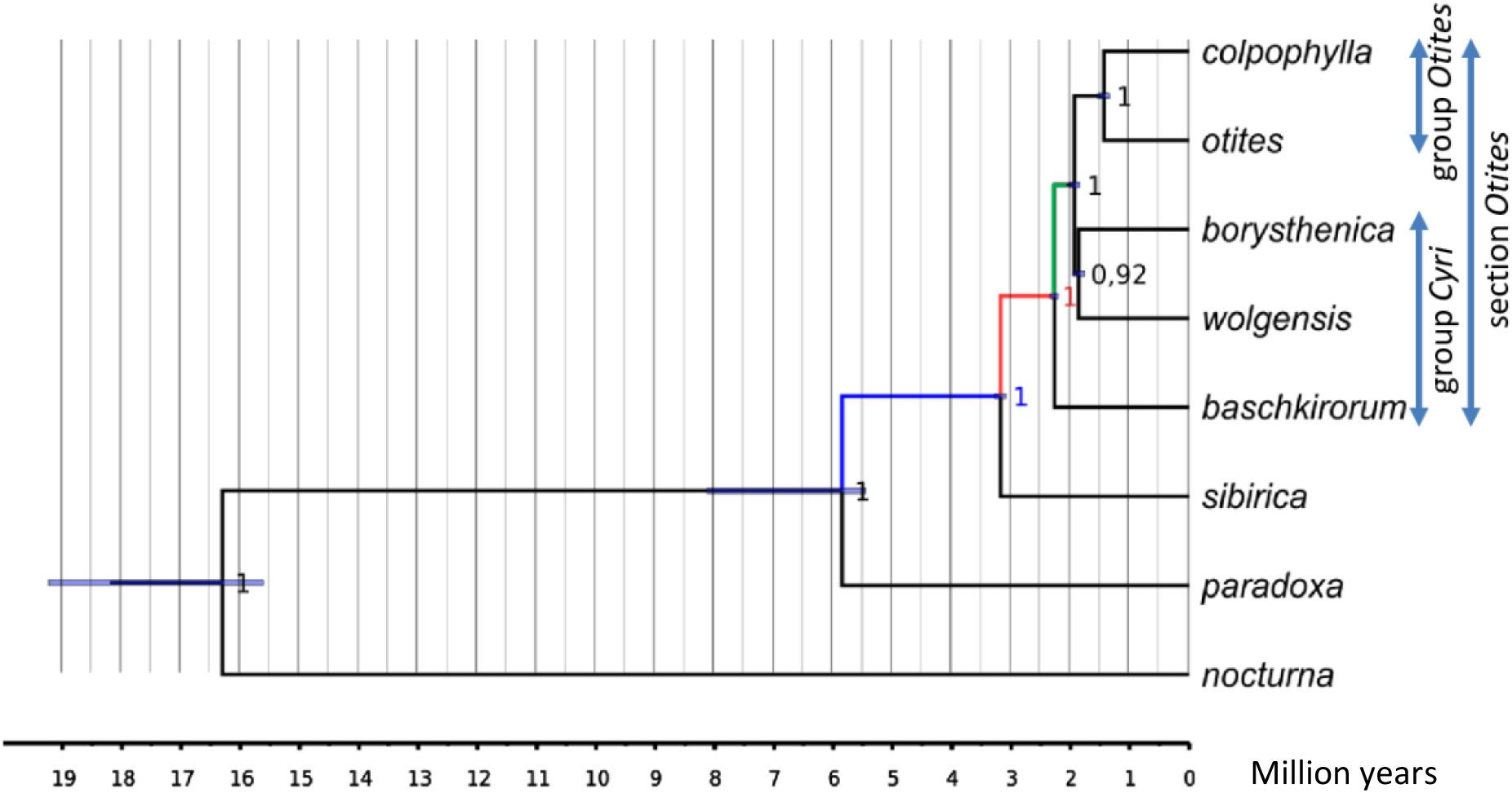
Phylogenetic tree of the section *Otites* and the closely related species. The colors of the highlighted branches refer to analyses of adaptive analyses. For details, see Results and Discussion.

The ω ratio (ω = dN/dS) is one of the most widely used statistical tests used to quantify selection pressures acting on protein-coding regions. This measure quantifies selection pressures by comparing the rate of substitutions at silent sites (dS), which are presumed neutral, to the rate of substitutions at non-silent sites (dN), which possibly experience selection. The comparison of the two- and three-ratios branch models in the CODEML program of PAML package (Yang, 2007) shows that the ω value of the branch preceding sex chromosome evolution (ω = 0.16; blue in Fig. 1) is lower than the ω value of the branch where dioecy and sex chromosomes evolved (ω = 0.29; red in Fig. 1). This difference is highly significant (P < 10^−99^, likelihood ratio test; LRT). After the formation of the dioecy (green in Fig. 1), the ω value significantly decreased (ω = 0.26; P < 10^−99^, LRT).

Because the increase of the ω values can be caused by the changes in the number of either positively or neutrally selected sites, we compared branch-site models of these branches. The results are summarized in Table 1. The branch where dioecy evolved showed a significant percentage of positively selected codons (P < 10^−99^, LRT). On the other hand, we did not detect codons under positive selection in the branch preceding dioecy (blue in Fig. 1) (P > 0.99, LRT). Moreover, the ω values of sites under purifying selection did not increase significantly (P = 0.95, Wilcoxon test). To confirm these results, we performed tests for relaxed selection using RELAX program (Wertheim, Murrell, Smith, Kosakovsky Pond, & Scheffler, 2015) of HyPhy package (Kosakovsky Pond et al., 2020; Pond, Frost, & Muse, 2005). Transition from the gynodioecy to the dioecy was connected with the selection intensification (P = 0.00; K = 1.55). On the other hand, when the branch involving dioecy evolution and the branch after sex chromosome evolution were compared, non-significant relaxation was detected (P = 0.339, K = 0.91). Most of the identified adaptively evolved codons (63 out of 87 codons) are recruited from the neutrally evolved codons, which is significantly more than by chance (P < 10^−16^, chi-squared goodness of fit test). Because the dioecy in the *Silene* section *Otites* most likely evolved from the gynodioecy, and because the gynodioecy is in the genus *Silene* usually of nucleo-cytoplasmic type, the dioecy evolved, in this case, most likely by the genetic fixation of a male sterile cytoplasm and subsequent recruitment of a fertility restorer as sex determining gene, as discussed by (Zluvova et al., 2005). In this case, no drastic change in population size or rate of inbreeding is necessary to open the route to dioecy. The observed absence of any sign of selective pressure relaxation is in accordance with this hypothesis.

**Table 1.**
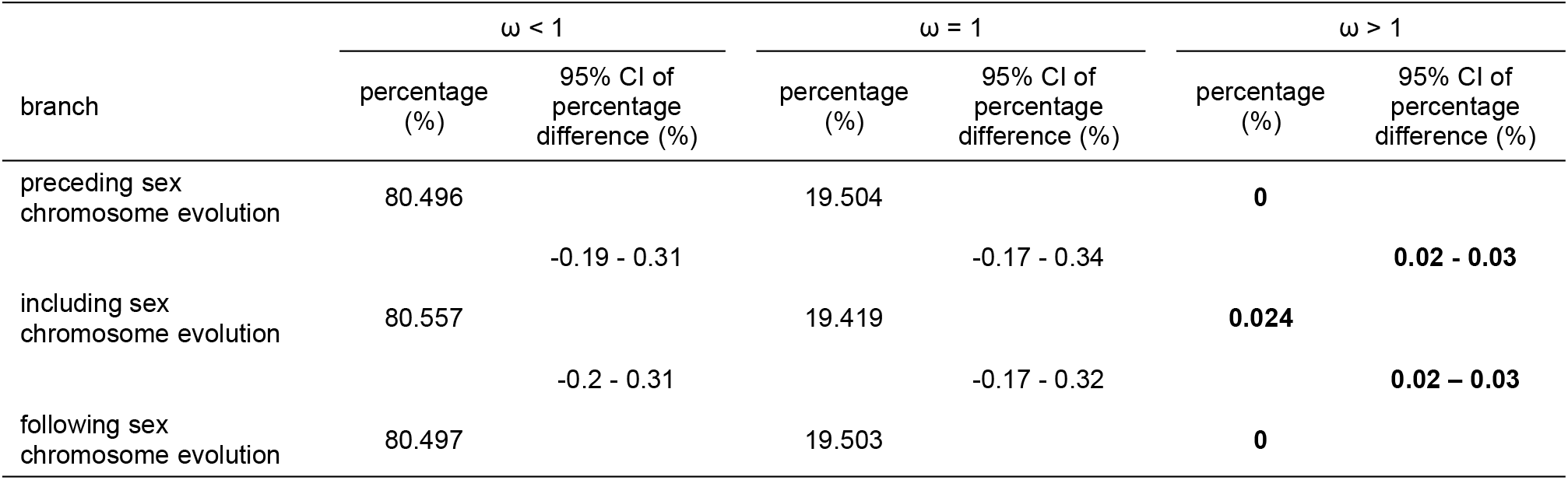
Branch-site analysis of *Silene* transcriptome before and after sex chromosome evolution. Confidence intervals (CI) are given for the difference of percentage between the neighboring rows. Statistically significant values are highlighted in bold.

| branch | $\omega < 1$ | | $\omega = 1$ | | $\omega > 1$ | |
| --- | --- | --- | --- | --- | --- | --- |
|  | percentage (%) | 95% CI of percentage difference (%) | percentage (%) | 95% CI of percentage difference (%) | percentage (%) | 95% CI of percentage difference (%) |
| preceding sex chromosome evolution | 80.496 |  | 19.504 |  | **0** |  |
|  |  | -0.19 - 0.31 |  | -0.17 - 0.34 |  | **0.02 - 0.03** |
| including sex chromosome evolution | 80.557 |  | 19.419 |  | **0.024** |  |
|  |  | -0.2 - 0.31 |  | -0.17 - 0.32 |  | **0.02 - 0.03** |
| following sex chromosome evolution | 80.497 |  | 19.503 |  | **0** |  |

The rise of the amount of positively selected codons on the branch including sex chromosome evolution cannot be caused by the selection of sexually antagonistic genes on the sex chromosomes (Rice, 1996), because the analysed dataset does not contain any completely sex-linked gene. Moreover, it cannot be caused by the accumulation of sexually antagonistic genes in the pseudoautosomal region, because we found that only one of the studied ESTs was located in this part of the sex chromosomes.

Because all close relatives of the analyzed species are gynodioecious, most likely with cytoplasmic male sterility, one plausible explanation for the rise of adaptive evolution can be connected with the presumed nucleo-cytoplasmic origin of the sex-determining system. The fixation of a certain type of male-sterility causing cytoplasm could influence the evolution of nuclear genes involved in mitochondrial metabolism. However, among the 42 ESTs identified as having at least one adaptively evolved codon on the branch involving sex chromosome formation, we did not find an overrepresentation of proteins located in mitochondria (7 ESTs; P = 0.77, Pearson’s chi-squared test).

We can hypothesize that this rise in the amount of positively selected codons is caused by the selection of genes involved in the specialization of the genome on the dioecy as described in the sex-allocation theory (Charnov, 1982), which states that female plants allocate more resources to the quality of seeds whereas male plants allocate resources into the amount of pollen. However, based on phenotypic data, it is possible to conclude that the adaptation of the genome to the dioecy is rather complex and includes a wide variety of genes (Geber, Dawson, & Delph, 1999). Indeed, the genes identified as adaptively evolving on the branch where dioecy evolved (see Supporting Information) show rather diverse characteristics. These results are in good accordance with the previous observations on the phenotypic level (Geber et al., 1999).

The discovered coincidence between the increased amount of adaptively evolved codons in autosomes and the evolution of dioecy cannot prove causality in this process. However, the hypothesis that the changes in autosomes represent an adaptation to the sexually dimorphic phenotype of dioecious species appears most likely.

## Materials and methods

The dataset from the previous work (Balounova et al., 2019) was supplemented by *S. nocturna* (SRR6040876) and *S. paradoxa* (SRR999296-SRR999299, publically available) samples. *S. pseudotites* has been excluded from the dataset based on its suspected hybrid origin. The reads of the *S. nocturna* and *S paradoxa* were assembled using Trinity (Haas et al., 2013). The assembly was treated with an Evigene pipeline to reduce the level of duplicates and used as a reference. The reads were mapped using Bowtie2 (Langmead & Salzberg, 2012), the SNPs were called via FreeBayes (Garrison & Marth, 2012) and phasing was performed with WhatsHap (Patterson et al., 2015). The regions with coverage lower than 2 masked using the maskfasta method in BEDTools (Quinlan & Hall, 2010). The phased sequences were added to the original alignments using the MAFFT aligner (Katoh & Standley, 2013) based on the results of best reciprocal blast hit search (Camacho et al., 2009). Phylogenetic analysis was done using the StarBeast2 module in BEAST 2 (Bouckaert et al., 2014; Ogilvie, Bouckaert, & Drummond, 2017). Two independent chains were run for 1000 000 000 states (Yule model, birth-death). For the dating, the calibration of the most recent common ancestor of *S. nocturna* and the other species was done similarly to Balounova et al. (Balounova et al., 2019) (based on estimated divergence per synonymous site dS (substitutions per synonymous site) and the divergence time estimates for several angiosperm lineages (Brassicaceae, Malvaceae, Euphorbiaceae, Fabaceae, Cucurbitaceae, Rosaceae, Solanaceae, and Poaceae), so as not to depend on a single fossil record or phylogenetic tree position, which yielded a mean substitution rate of 5.35 × 10^−9^ synonymous substitutions/site/ year) (De La Torre, Li, Van de Peer, & Ingvarsson, 2017). Ks values for the distance of individual species of the section *Otites* to *S. nocturna* were estimated using KaKs calculator 2 program (Wang, Zhang, Zhang, Zhu, & Yu, 2010), and the mean value was used for the calibration of the tree. The resulting dataset did not contain any of the completely sex-linked genes identified previously (Balounova et al., 2019) (Martin et al., 2019).

PAML analyses were used to determine whether some ESTs evolved under selective pressure. The CODEML program of PAML (Yang, 2007) was used to estimate the ratio (ω) of the non-synonymous substitution rate (dN) to the synonymous substitution rate (dS). As the reference tree, the phylogenetic tree constructed as described above was used. The equilibrium frequencies of codons were calculated from the nucleotide frequencies (CodonFreq = 2) because it best fits the data as calculated by second-order AIC. All models in CODEML were run at five different initial ω values (ω = 0.1, 0.2, 0.5, 1, 2), and no problems with the convergence were observed. Both branch and branch-site models were applied to the branch preceding sex chromosome formation, to the branch including sex chromosome formation and to the branch following the sex chromosome formation. In the branch analyses, two-ratios models were compared to three-ratios models to reveal whether the ω values differ significantly. The resulting log likelihood values were evaluated using likelihood-ratio tests to determine any statistical significance of the difference. The chi2 program of PAML was used to estimate the P-values. The confidence intervals for proportion were computed online (http://vassarstats.net/prop1.html) using the method with continuity correction (Newcombe, 1998) that is derived from a procedure outlined in (Wilson, 1927). The results obtained in CODEML were further confirmed using a test for relaxed selection in RELAX program (Wertheim et al., 2015) of the HypHy package (Pond et al., 2005) (Kosakovsky Pond et al., 2020). Because the results did not converge well, the analysis was run 500-times and the results with the highest log likelihood were used.

The modified model A of the branch-site models was used to compute the percentages of purifying, neutral and adaptive codons in the analyzed ESTs, to compute the ω values, to compute the Bayes Empirical Bayes probability of codons being either purifying, neutral or adaptive and to identify ESTs with adaptively evolved codons. The Wilcoxon test computed online (https://www.socscistatistics.com/tests/signedranks/default2.aspx) was used to compare the ω values of the codons with ω < 1 between the branches before and during sex chromosome and dioecy formation. The chi-squared goodness of fit test was used to calculate the P-value of adaptively evolving codons to evolve from neutrally evolving codons. The Pearson’s chi-squared test was used to compute the P-value of a random presence of genes acting in mitochondria among the genes with positively selected codons.

## Acknowledgments

This research was supported by the Czech Science Foundation (grant 19-15609S). Computational resources were supplied by the project “e-Infrastruktura CZ” (e-INFRA LM2018140) provided within the program Projects of Large Research, Development and Innovations Infrastructures and by the ELIXIR-CZ project (LM2018131), part of the international ELIXIR infrastructure.

## Competing Interest Statement

No financial and/or non-financial competing interests declared.

## Notes

**Competing Interest Statement:** The authors declare no competing interest.

### Competing Interest Statement

The authors have declared no competing interest.

